# E(3) equivariant graph neural networks for robust and accurate protein–protein interaction site prediction

**DOI:** 10.1101/2022.12.14.520476

**Authors:** Rahmatullah Roche, Bernard Moussad, Md Hossain Shuvo, Debswapna Bhattacharya

## Abstract

Artificial intelligence-powered protein structure prediction methods have led to a paradigm-shift in computational structural biology, yet contemporary approaches for predicting the interfacial residues (i.e., sites) of protein-protein interaction (PPI) still rely on experimental structures. Recent studies have demonstrated benefits of employing graph convolution for PPI site prediction, but ignore symmetries naturally occurring in 3-dimensional space and act only on experimental coordinates. Here we present EquiPPIS, an E(3) equivariant graph neural network approach for PPI site prediction. EquiPPIS employs symmetry-aware graph convolutions that transform equivariantly with translation, rotation, and reflection in 3D space, providing richer representations for molecular data compared to invariant convolutions. EquiPPIS substantially outperforms state-of-the-art approaches based on the same experimental input, and exhibits remarkable robustness by attaining better accuracy with predicted structural models from AlphaFold2 than what existing methods can achieve even with experimental structures. Freely available at https://github.com/Bhattacharya-Lab/EquiPPIS, EquiPPIS enables accurate PPI site prediction at scale.

## Introduction

Protein-protein interactions (PPI) underpin numerous biological processes^1,2^. Despite their importance, experimental characterization of PPI remains challenging due to the costly and time-consuming nature of the experimental assays^3^. Computational methods offer a cheaper and high-throughput alternative by predicting the bound complex structures of interacting proteins from the sequences and/or the unbound structures of individual protein chains. A closely related problem—and the one addressed in this study—is the prediction of the PPI sites, which are the interfacial residues of the interacting protein chains.

Accurately predicting the interface of interacting proteins and the identification of the PPI sites remain challenging even after decades of research^4,5^. Various methods have been proposed, but with limited success. Partner-independent PPI site prediction^6–11^, which involves the prediction of putative interaction sites based only upon the surface of an isolated protein, without any knowledge of the partner or complex, is even more challenging compared to partner-aware PPI site prediction^12–16^ due to the absence of any information about the partner protein and auxiliary information on the complex interfaces. In this work, we focus on partner-independent PPI site prediction.

Predicting how proteins interact, and in particular, predicting the PPI sites, has a long history ^11,14,17–24^. While initial models focused on feature engineering with machine learning^6,18,25,26^, subsequent work sought to capture more complex patterns using deep learning ^7,9,12,13,16^. The vast majority of the existing methods rely on readily available protein sequence information, but their predictive accuracies are often quite limited^27^. Structure-based methods that integrate known structural information from the Protein Data Bank (PDB^28^) are usually more accurate. However, these approaches are limited by the paucity of experimentally solved protein structures in the PDB. In the 14th edition of the Critical Assessment of Structure Prediction (CASP14) experiment, AlphaFold2^29^ attained an unprecedented performance level, enabling highly accurate prediction of single-chain protein structural models at proteome-wide scale^30,31^. Given the recent progress, a natural question arises: can we leverage the predicted structural information by AlphaFold2 for accurate partner-independent PPI site prediction at scale?

In the recent past, representation learning with graph structured data has been prevailing in different applications. In particular, graph neural networks (GNNs) have surged as the major choice for deep graph learning^32–34^. GNNs are permutation equivariant networks that operate on graph structured data, with numerous applications ranging from dynamical systems to conformational energy estimation^35,36^. However, off-the-shelf GNNs do not take into account symmetries naturally occurring in 3-dimensional space. That is, they ignore the effects of invariance and equivariance with respect to the E(3) symmetry group, i.e., the group of rotations, reflections, and translations in 3D space. The recent E(n) equivariant graph neural networks^37^ address this problem by being translation, rotation, and reflection equivariant in 3D space that can be scaled to higher dimensional spaces (E(n)), while preserving permutation equivariance. SE(3) equivariant neural networks^38^ are another recent graph-based models that can deal with the absolute coordinate systems in 3D space, but SE(3) equivariant models do not commute with reflections of the input. E(3) equivariant neural networks, on the other hand, transform equivariantly with translation, rotation and reflections, which make them suitable for molecular data where chirality of the molecules are often important, such as proteins, particularly when predicted protein structures are used as input that may contain mirror images. As such, E(3) equivariant graph neural networks offer an elegant choice for partner-independent PPI site prediction, where the input consists of a 3D structure of an isolated protein, known to be involved in PPIs, but where the structure of the partner or complex is not known. Being designed from geometric first-principles, symmetry-aware models such as E(3) equivariant graph neural networks are highly suitable for 3D molecular data, providing richer representation while avoiding expensive data augmentation strategies.

The contribution of the present work is the introduction of a symmetry-aware PPI site prediction method, EquiPPIS, built on E(3) equivariant graph neural networks that yields state-of-the-art accuracy by substantially outperforming existing approaches based on the same experimental input. What is more striking is that EquiPPIS attains better accuracy with AlphaFold2-predicted structural models as input than what existing methods can achieve even with experimental input. We directly verify that the performance gains are connected to the unique E(3) equivariant architecture of EquiPPIS. The robustness and performance resilience of our method enable large-scale PPI site prediction without compromising on accuracy.

## Results

E(3) equivariant graph neural networks for protein–protein interaction site prediction

**Fig. 1** illustrates our E(3) equivariant graph neural network model for partner-independent PPI site prediction. Different from the recent structure-aware graph learning approaches for PPI site prediction^9^ that only exploit pairwise distances between all residue pairs (i.e., distance maps) as the spatial information, our E(3) equivariant graph neural network model directly leverages the C_α_ coordinates extracted from the input monomer together with sequence- and structure-based node and edge features. By using an E(3) equivariant architecture, our symmetry-aware model can learn to preserve the known transformation properties of 3D coordinates under translation, rotation, and reflection, improving PPI site prediction. The EquiPPIS method consists of three major modules. The first module (**Fig. 1a**) converts the input protein monomer into an undirected graph 𝒢 = (𝒱,ℰ), with 𝒱 denoting the residues (nodes) and ℰ denoting the interaction between nonsequential residue pairs according to their pairwise spatial proximity (edges). The spatial proximity between nonsequential residue pairs (i.e., having sequence separation greater or equal to 6) is determined by calculating the Euclidean distances between the C_α_ atom of all residue pairs and then setting a cutoff distance of 14Å to obtain the interacting pairs. The sequence- and structure-based node and edge features (see the Methods section) are then fed into the second module together with the C_α_ coordinates extracted from the input monomer. The second module (**Fig. 1b**) is a deep E(3) equivariant graph neural network that conducts a series of transformations of its input through a stack of equivariant graph convolution layer (EGCL)^37^, each updating the coordinate and node embeddings using the edge information and the coordinate and node embeddings from the previous layer. Finally, a sigmoidal function is applied to the last EGCL node embedding to predict the probability of every residue in the input monomer to be a PPI site, thereby converting the PPI site prediction into a graph node classification task (**Fig. 1c**).

**Fig. 1:**
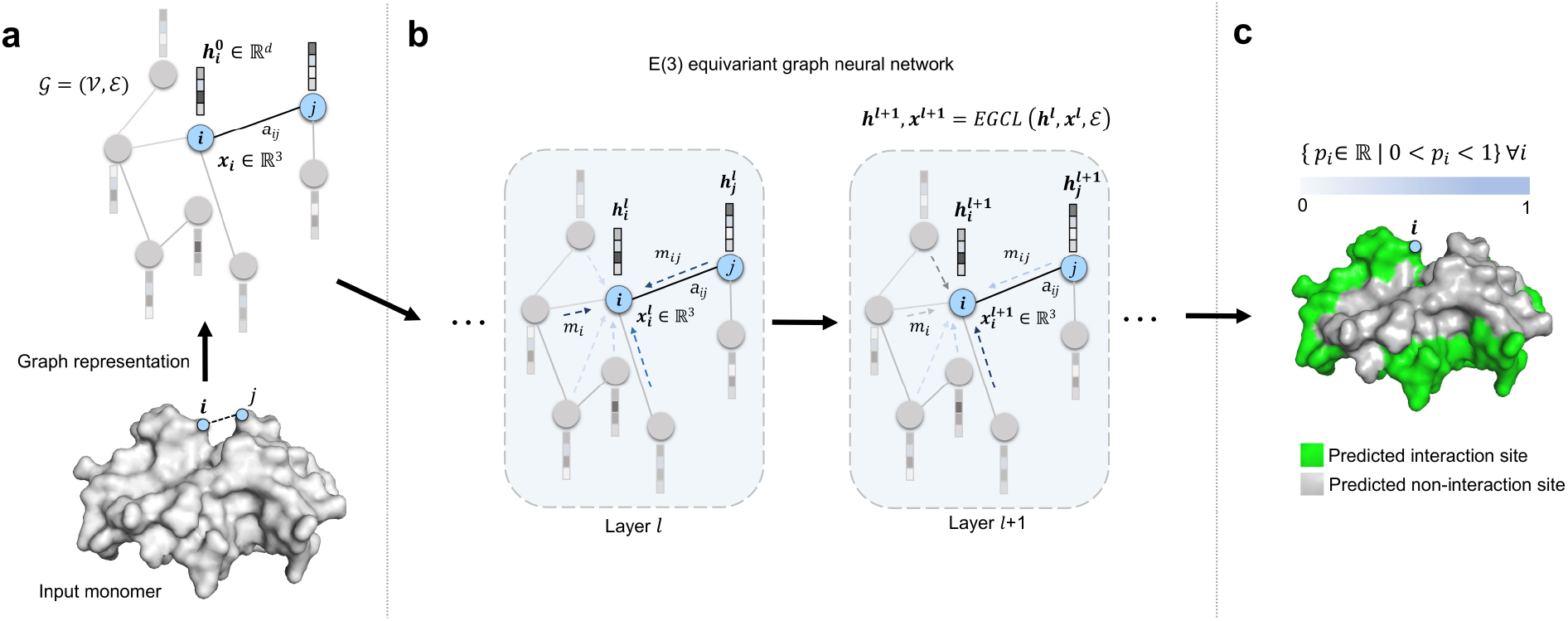
Illustration of the EquiPPIS method for protein–protein interaction site prediction. (**a**) The input protein monomer is converted into an undirected graph. (**b**) Equivariant graph convolutions are then employed on the input graph. (**c**) PPI sites are finally predicted as a graph node classification task.

### Experiments

For training and performance evaluation, we use a combination of three widely used and publicly available benchmark datasets: Dset_186^6^, Dset_72^6^, and Dset_164^39^, named by the number of proteins in the datasets. We follow the same train and test splits from the recent PPI site prediction method GraphPPIS^9^, which combines the aforementioned three datasets and subsequently removes redundancy, resulting in 335 targets as the training set (Train_335) and 60 targets as the test set (Test_60). Details of our training procedure are provided in the Methods section. We evaluate our proposed method on a diverse series of challenging test scenarios using standard performance evaluation metrics including accuracy, precision, recall, F1-score (F1), Matthews correlation coefficient (MCC), area under the receiver operating characteristic curve (ROC-AUC), and area under the precision-recall curve (PR-AUC). First, we demonstrate that EquiPPIS improves upon state-of-the-art accuracy on Test_60 dataset by comparing it directly against a wide variety of existing approaches including PSIVER^6^ (based on Naïve Bayes), ProNA2020^40^ (based on neural network), SCRIBER^41^ (based on two layers of logistic regression), DLPred^42^ (based on long short-term memory), DELPHI^8^ (based on an ensemble of convolutional and recurrent neural networks), DeepPPISP^7^ (based on convolutional neural network), SPPIDER^10^ (based on support vector machine, neural network, and linear discriminant analysis), MaSIF-site^43^ (based on geometric deep learning), and GraphPPIS (based on deep graph convolutional network). Among the competing methods, PSIVER, ProNA2020, SCRIBER, DLPred, and DELPHI are purely sequence-based methods; whereas DeepPPIS, SPPIDER, MaSIF-site, GraphPPIS, and our new method EquiPPIS additionally integrate structural information. Next, we examine the reasons for such high performance attained by EquiPPIS and verify that it is indeed connected to the equivariant nature of the model used. To broaden the applicability of our method beyond predicting PPI sites based only on experimentally-solved input monomers extracted from the bound complex structures, we explore a number of extensions including using unbound experimental structures and computationally predicted structural models as input. Specifically, using a subset of 31 proteins from the Test_60 dataset with known unbound monomeric structures in the PDB as an additional unbound test set (hereafter called UBtest_31), we show that EquiPPIS exhibits remarkable robustness and performance resilience compared to the existing approaches. Furthermore, by replacing the experimental input monomers with AlphaFold2 predicted structural models for the Test_60 dataset, we demonstrate that EquiPPIS attains state-of-the-art accuracy, which is better than what the top performing competing method can achieve even with experimental structures. The superior performance of EquiPPIS even when using predicted structural models as input dramatically enhances the scalability of partner-independent PPI site prediction without compromising on accuracy. Finally, we examine the relative importance of each feature we adopted by conducting feature ablation experiments using an independent validation set consisting of 42 targets (hereafter called Validation_42) collected from the Test_315 dataset of the published work of GraphPPIS after filtering out proteins with >25% pairwise sequence identity with our test sets. We also use this validation set for hyperparameter selection.

### Test set performance

We compare EquiPPIS with five sequence-based (PSIVER, ProNA2020, SCRIBER, DLPred and DELPHI) and four structure-based (DeepPPISP, SPPIDER, MaSIF-site, and GraphPPIS) predictors on the Test_60 set. As shown in **Table 1**, in addition to outperforming the sequence-based methods (PR-AUC ranging from 0.190 to 0.319) by a large margin, EquiPPIS significantly improves upon state-of-the-art accuracy by outperforming the structure-based methods. Remarkably, EquiPPIS is the only method attaining ROC-AUC of more than 0.8, which is noticeably better than the closest competing method GraphPPIS. Interestingly, the published work of GraphPPIS sets the goal of achieving ROC-AUC of 0.8 as a motivation for future work, while acknowledging it as one of the current impediments. In summary, EquiPPIS is a leap forward for partner independent PPI site prediction.

**Table 1.**
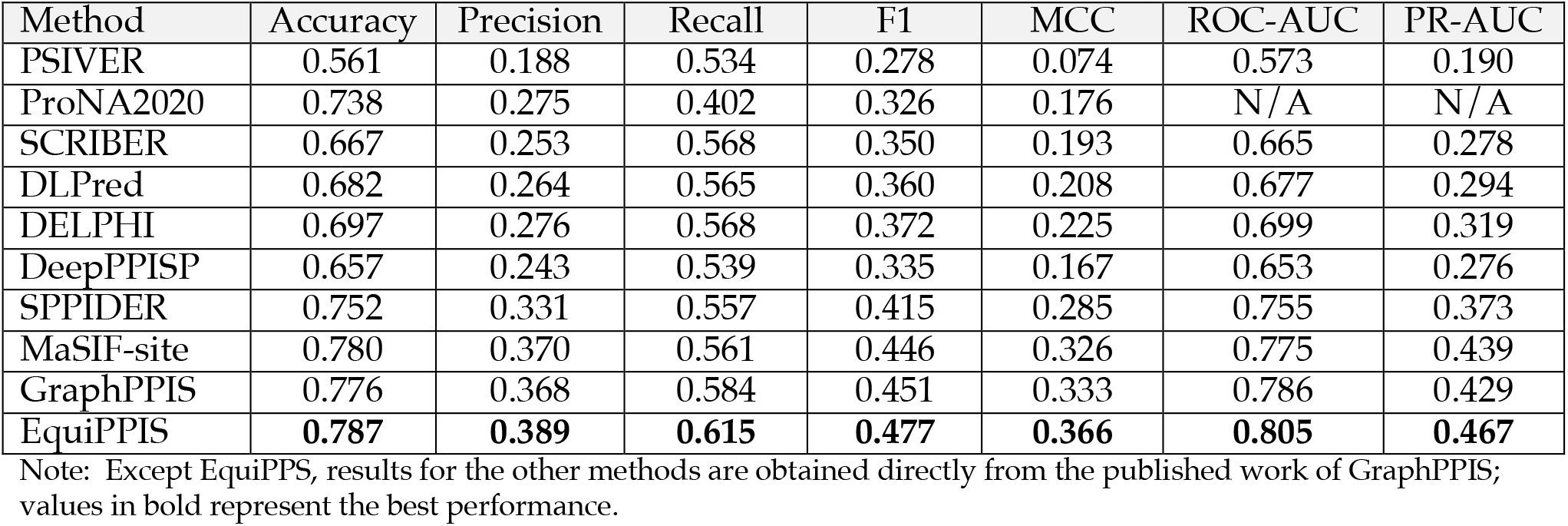
PPI site prediction performance on the Test_60 dataset for various methods.

**Fig. 2** presents two representative examples from the Test_60 dataset comparing the PPI site predictions using EquiPPIS and GraphPPIS. For the first example of a sugar binding protein of Trichosanthes kirilowii (PDB ID: 1GGP, chain A) having length 234 (**Fig. 2a**), EquiPPIS correctly predicts majority of the observed PPI sites, attaining Precision, Recall, F1, and MCC of 0.8, 0.545, 0.649, and 0.601, respectively; whereas GraphPPIS fails to predict any correct PPI sites with Precision, Recall, F1 and MCC of 0, 0, 0, -0.185, respectively. The second example is a Hydrolase inhibitor of Triticum aestivum in complex with Bacillus subtilis (PDB ID: 2B42, chain A) having length 364 (**Fig. 2b**), where GraphPPIS predicts many false positive PPI sites, resulting in low Precision, Recall, F1 and MCC of 0.105, 0.231, 0.144, and -0.004, respectively. EquiPPIS on the other hand attains reasonably accurate predictive performance having Precision, Recall, F1, and MCC of 0.595, 0.564, 0.579, and 0.53, respectively. In both cases, EquiPPIS predictions are strikingly similar to the experimentally observed PPI sites.

**Fig. 2:**
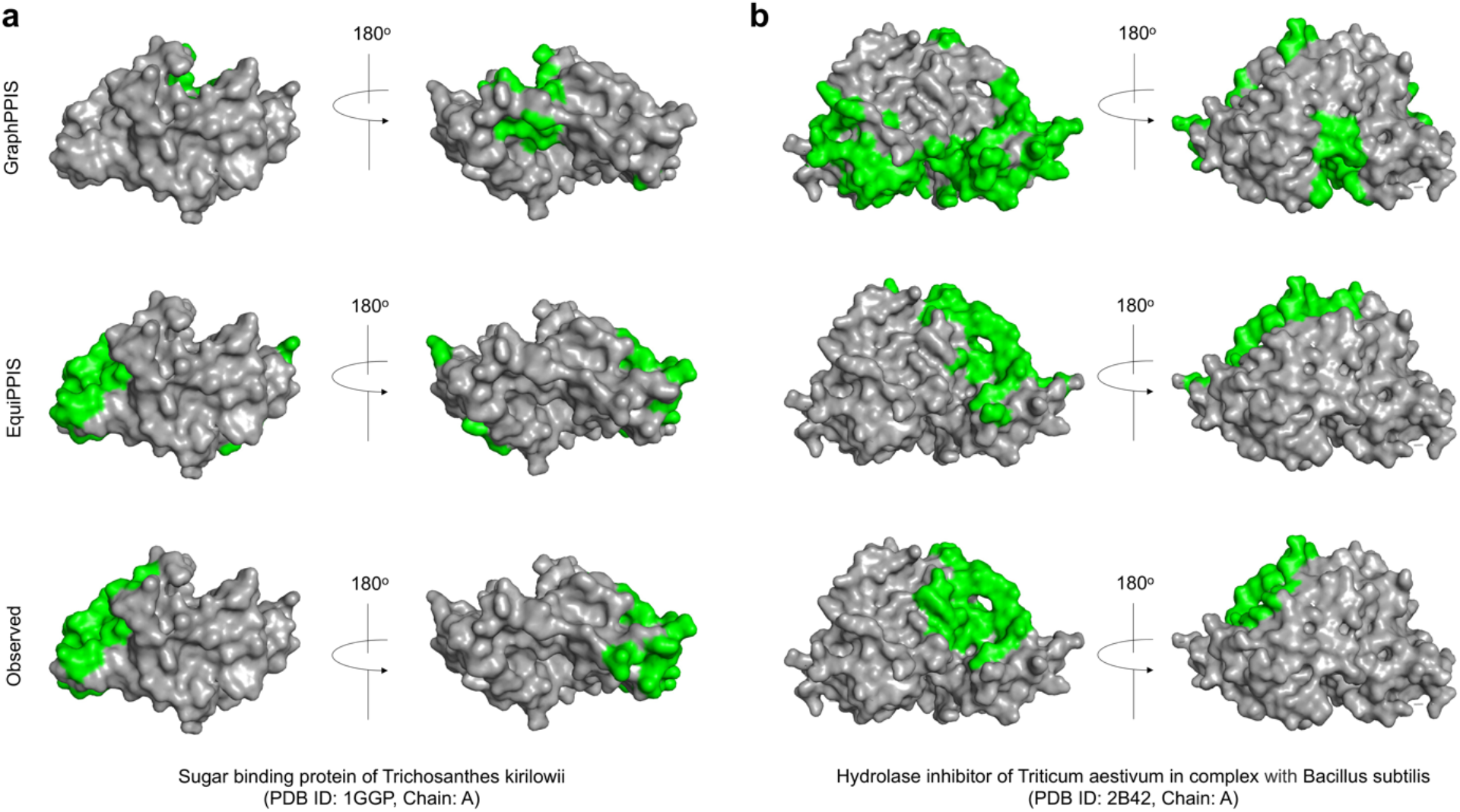
GraphPPIS and EquiPPIS predictions compared to the experimental observation. (**a**) Sugar binding protein of Trichosanthes kirilowii. (**b**) Hydrolase inhibitor of Triticum aestivum in complex with Bacillus subtilis. The regions highlighted in green represent PPI sites.

### Analyzing the importance of equivariance

In the above experiments, EquiPPIS exhibits significantly improved performance. In order to gain insight into the reasons behind such high performance and verify that it is connected to the equivariant nature of the model, we perform a series of experiments by gradually isolating the effect of the equivariant graph convolutions used in EquiPPIS. In particular, we train several baseline models and compare them head-to-head with the full-fledged version of EquiPPIS. First, we train a baseline network by turning off the coordinate updates of the equivariant graph convolution layers, thus making it an invariant network (hereafter called ‘EquiPPIS invariant’). Since the full-fledged version of EquiPPIS employs attention operations for aggregated embedding as part of the equivariant message passing, we train another baseline network where attention operation is turned off during equivariant message passing, resulting in an equivariant network but without attention (hereafter called ‘EquiPPIS w/o attention’). Additionally, we train two off-the-shelf GNNs for PPI site prediction: graph convolution network (GCN)^34^ and graph attention network (GAT)^44^. All baseline networks are trained on the same Train_335 dataset using the same set of input features and hyperparameters as the full-fledged version of EquiPPIS (see the Methods section). **Fig. 3a-3d** shows the performance of EquiPPIS compared to the baseline networks on the Test_60 set. The results demonstrate that the full-fledged version of EquiPPIS outperforms all baseline models. For example, we observe that the full-fledged version of EquiPPIS attains an ROC-AUC of more than 0.8, which is the best accuracy compared to all baseline models. The ‘EquiPPIS invariant’ baseline, however, falls short of achieving an ROC-AUC of 0.8, suggesting that it is the equivariant nature of EquiPPIS that is responsible for the accuracy gain. Turning off the attention operation as done in the ‘EquiPPIS w/o attention’ baseline leads to an accuracy decline (an ROC-AUC of 0.8) compared to the full-fledged version of EquiPPIS, but still better than the invariant network. That is, attention operation during equivariant message passing contributes to an improvement in accuracy. It is worth noting that despite the accuracy drop from the full-fledged version of EquiPPIS, both ‘EquiPPIS invariant’ and ‘EquiPPIS w/o attention’ baselines outperform GraphPPIS. On the other hand, off-the-shelf GCN- and GAT-based baselines exhibit much lower accuracies compared to GraphPPIS, let alone EquiPPIS. Overall, the results underscore the importance of equivariance in particular and symmetry-aware nature of the new EquiPPIS model in general for improved predictive accuracy.

**Fig. 3:**
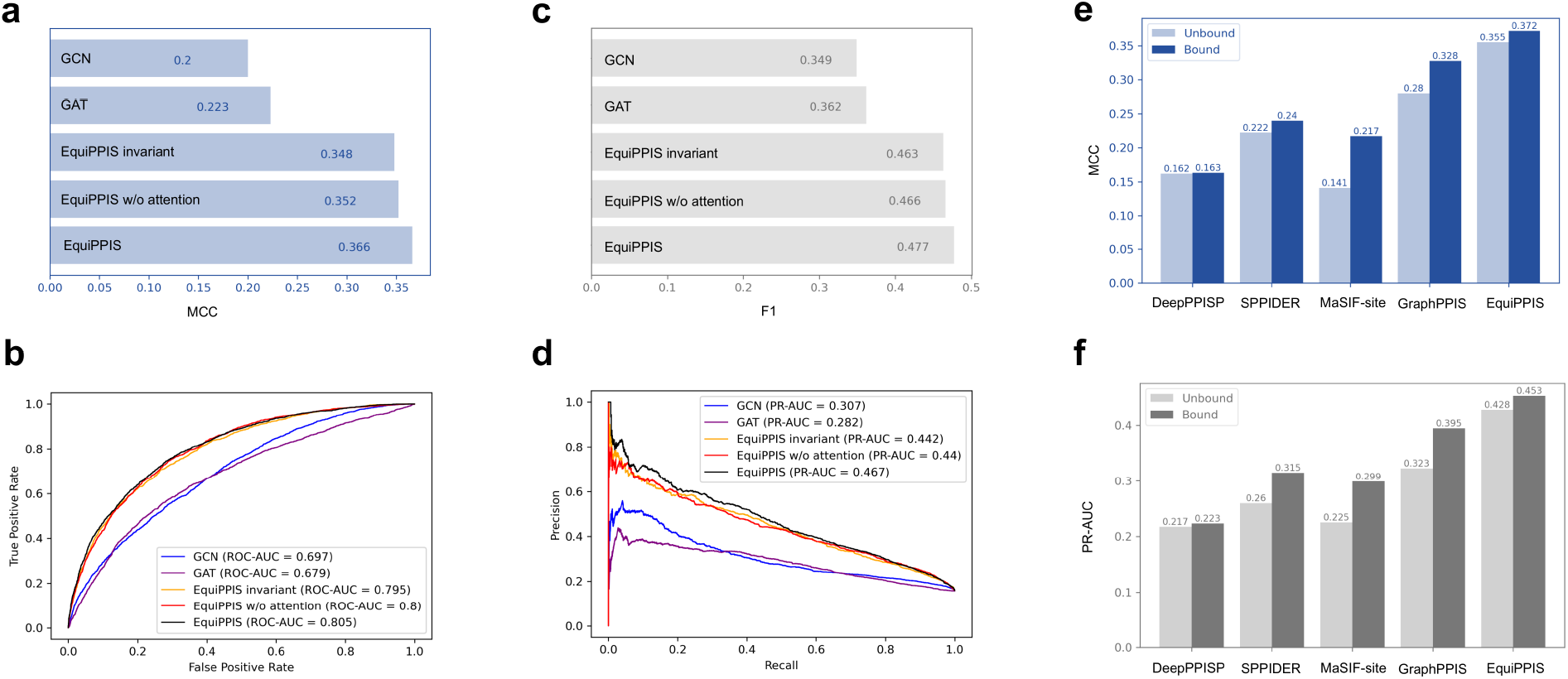
Performance analysis on Test_60 set (**a**-**d**) and on UBtest_31 set (**e**-**f**). (**a**) MCC, (**b**) ROC-AUC, (**c**) F1, and (**d**) PR-AUC of EquiPPIS on Test_60 set compared to the baseline models ‘EquiPPIS invariant’, ‘EquiPPIS w/o attention’, graph convolution network (GCN), and graph attention network (GAT). (**e**) MCC and (**f**) PR-AUC of EquiPPIS on UBtest_31 set compared to other structure-based methods GraphPPIS, MaSIF-site, SPPIDER, and DeepPPISP.

In addition to prediction accuracy, robustness of model is another key aspect to consider. While experimentally-solved bound complex structures are used during EquiPPIS training, protein– protein binding often leads to conformational changes by “induced fit” mechanism (binding first) or “conformational selection” (conformational change first)^45^. To evaluate the robustness of EquiPPIS and the effect of conformational changes, we examine the impact on the accuracy when unbound structures are used during prediction instead of their bound states for EquiPPIS as well as the other structure-based PPI site predictors (DeepPPISP, SPPIDER, MaSIF-site, and GraphPPIS) using the unbound test set (UBtest_31) of 31 proteins. As shown in **Fig. 3e-3f**, EquiPPIS outperforms all other methods by a large margin, while having the least impact on accuracy when unbound structures are used during prediction. For example, the closest competing methods MaSIF-site and GraphPPIS suffer from significant accuracy drop both in terms of MCC (35%, and 14.6% drop, respectively) and PR-AUC (24.7%, and 18.2% drop, respectively), whereas EquiPPIS experiences only 4.6%, and 5.5% drop in MCC and PR-AUC, respectively. What is most striking is that the accuracy gap between EquiPPIS and the competing methods is so large that EquiPPIS using unbound structures attains much better accuracy even when the competing methods are using the bound structures. That is, EquiPPIS exhibits remarkable robustness and performance resilience compared to existing approaches.

### Beyond experimental input: state-of-the-art performance with AlphaFold2

EquiPPIS achieves state-of-the-art accuracy with experimental structures as input in both bound and unbound states. A natural question to ask is can we achieve similar predictive accuracy when computationally predicted structural models are used as input instead of experimental structures? Given the exceptional performance of AlphaFold2 in the CASP14 experiment and the open availability of the AlphaFold2 protocol, it is now possible to predict single-chain protein structural models from the amino acid sequence with high degree of accuracy. In principle, a robust method such as EquiPPIS should be able to generalize when predicted structural models are used without significant drop in accuracy, thereby broadening its applicability beyond experimental input. Motivated by the prospect, we examine the impact on the accuracy by replacing the experimental input with AlphaFold2 predicted structural models for the Test_60 dataset. **Table. 2** shows the performance of EquiPPIS compared to the closest competing method GraphPPIS. Remarkably, EquiPPIS using AlphaFold2-predicted structural models attains better accuracy (PR-AUC = 0.451) than GraphPPIS using experimental structures (PR-AUC = 0.429), let alone GraphPPIS using predicted structural models (PR-AUC = 0.399). While there is a performance decline for both methods when switching from experimental input to prediction, the performance drop for EquiPPIS is lower (ΔPR-AUC = 0.016) than that of GraphPPIS (ΔPR-AUC = 0.03). The results demonstrate the generalizability of EquiPPIS, thus opening the possibility of large-scale PPI site prediction by utilizing high-throughput computational prediction without compromising on accuracy.

**Table 2.** Performance comparison between EquiPPIS and GraphPPIS with experimental input and AlphaFold2 predicted structural models for the Test_60 dataset.

| Method | Input type | Accuracy | Precision | Recall | F1 | MCC | ROC-AUC | PR-AUC |
| --- | --- | --- | --- | --- | --- | --- | --- | --- |
| EquiPPIS | Experimental | **0.787** | **0.389** | **0.615** | **0.477** | **0.366** | **0.805** | **0.467** |
|  | AlphaFold2 | 0.780 | 0.379 | 0.615 | 0.469 | 0.356 | 0.795 | 0.451 |
| GraphPPIS | Experimental* | 0.776 | 0.368 | 0.584 | 0.451 | 0.333 | 0.786 | 0.429 |
|  | AlphaFold2 | 0.767 | 0.357 | 0.590 | 0.445 | 0.324 | 0.772 | 0.399 |

\* Results obtained directly from the published work of GraphPPIS; values in bold represent the best performance.

### Ablation studies and choice of hyperparameters

To examine the relative importance of the features adopted in EquiPPIS, we conduct feature ablation experiments by gradually isolating the contribution of individual feature or groups of features during model training and evaluating the accuracy on the independent validation set Validation_42. **Fig. 4a** shows the accuracy decline measured in terms of ΔPR-AUC and ΔROC-AUC when various features are isolated from the full-fledged version of EquiPPIS. The results demonstrate that all features contribute to the overall accuracy achieved by EquiPPIS. For example, we notice accuracy decline when we isolate the sequence-based features one by one including amino acid residue type (No AA), position specific scoring matrix (No PSSM), and protein language model ESM2 (No ESM2). Not surprisingly, the evolutionarily feature PSSM and protein language model-based ESM2 feature contribute more than just the residue type features. We notice a significant performance drop when all three sequence-based features are isolated (No seq). Similarly, we notice consistent accuracy decline when we discard the structure-based features individually including secondary structure (No ss), relative solvent accessibility (No rsa), local geometry (No local geom), residue orientation (No orient), contact count (No contact count) as well as relative residue positioning and residue virtual surface area (No res pos + area). Because we use multi-level discretization of secondary structure (e.g., 3-state and 8-state) and relative solvent accessibility (e.g., 2-state and 8-state) as well as backbone torsion angles, which are closely related to the secondary structure, we conduct feature ablation experiments by discarding the 8-state secondary structure, 8-state relative solvent accessibility, and backbone torsion angles. The resulting model (No multi-level) shows accuracy decline compared to the full-fledged version of EquiPPIS, indicating the effectiveness of combining multi-granular information. Finally, we also notice an accuracy drop when we isolate the edge feature (No edge) that takes into account the contributions of sequence separation and spatial interaction.

**Fig. 4:**
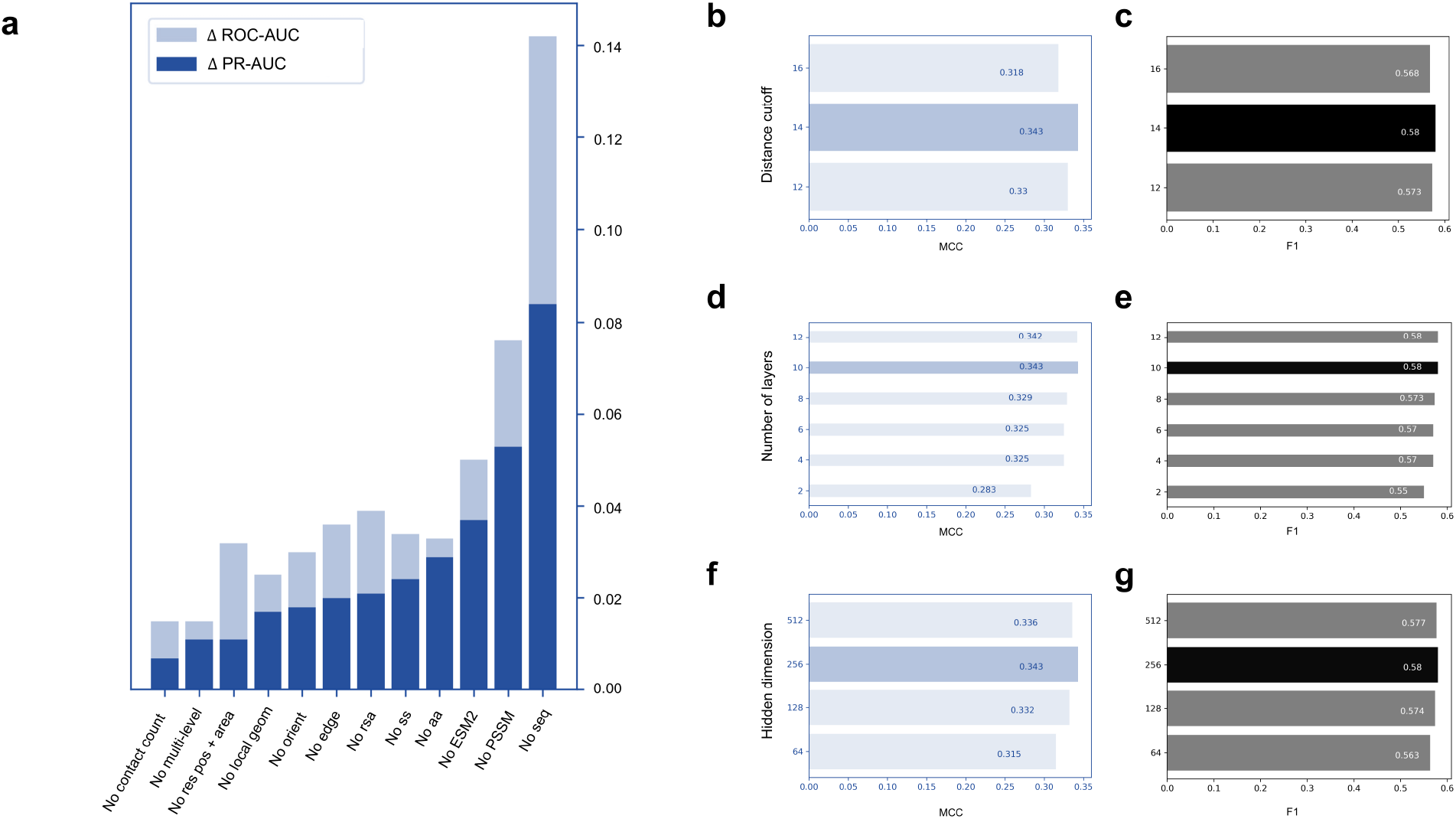
Ablation studies and hyperparameter selection. (**a**) Validation set accuracy decline after feature ablation. Dark and light blue bars indicate the PR-AUC decline (ΔPR-AUC) and the ROC-AUC decline (ΔROC-AUC), respectively. Validation set accuracy in terms of MCC and F1 for various choices of hyperparameters including (**b, c**) distance cutoff, (**d, e**) number of layers, and (**f, g**) hidden dimension. The selected hyperparameters yielding the best accuracy are highlighted in darker shade.

We also use the Validation_42 set to select the hyperparameters. Based on the results of the grid search as shown in Fig. **4b-4g**, we select a 10-layer EGCL framework with 256 hidden units and the cutoff distance used to obtain the interacting residue pairs is set to 14Å. We use the hyperparameters selected in the independent validation set during training and testing.

## Discussion

This work introduces EquiPPIS, a symmetry-aware deep graph learning model for protein– protein interaction site prediction based on E(3) equivariant graph neural networks. We demonstrate that EquiPPIS outperforms existing methods and despite being trained on experimental structures, it generalizes extremely well to predicted structural models from AlphaFold2 to the extent that EquiPPIS attains better accuracy with predicted structural models than what existing approaches can achieve even with experimental structures. Through controlled experiments, we verify the importance of equivariance as one of the major driving forces behind the improved performance. In addition to questions around the effect of equivariance on accuracy, our ablation study on an independent validation set confirms the contribution of various features adopted in EquiPPIS. Our study leads to a series of interesting questions to consider: of particular interest is the possibility of broadening the applicability of our method beyond experimental input for large-scale PPI site predictions with high accuracy by utilizing rapid computational prediction. Further, a promising direction for future work is to investigate the potential benefits of explicitly including multiple sequence alignment (MSA) information and measure the extent to which it may influence the accuracy. While an MSA-free method such as EquiPPIS offers some unique advantages by being broadly applicable even for proteins that do not have homologous sequences in the current sequence databases and bypasses the computational overhead of MSA searching, MSA may still provide a rich source of additional information for further improving the accuracy of PPI site prediction that might be worth exploring. Finally, while we find that EquiPPIS exhibits excellent predictive accuracy and remarkable robustness, an open challenge that remains is the interpretability of our deep learning model. The evolutionary and functional significance of the residues predicted to be in PPI site by means of the latent representation underlying the neural architecture of EquiPPIS still need to be systematically explored. We expect our proposed method can be easily extended to other biomolecular interaction site prediction tasks, including predicting protein-binding sites with other molecules, such as DNA, RNA and small ligands, with improved accuracy and robustness.

## Methods

### Graph representation and featurization

#### Graph representation

We represent the input protein monomer as a graph 𝒢 = (𝒱,ℰ), in which a node *v*∈ 𝒱 represents a residue and an edge *e*∈ ℰ represents an interacting residue pair. We consider two residues to be interacting if their C_α_ Euclidean distance is no more than 14Å. The cutoff 14Å is chosen on an independent validation set as presented in **Fig. 4**. To focus on longer-rage interactions, we only consider interacting residue pairs having a minimum sequence separation of 6.

#### Node features

We use three types of sequence-based node features: (1) one-hot encoding of the residue (i.e., a binary vector of 20 entries indicating its amino acid type), resulting in L×20 feature set, where L is the length of the input monomer; (2) position specific scoring matrix (PSSM) obtained by running PSI-BLAST^46^ to obtain L×20 feature set by considering the first 20 columns from the PSSM and normalizing the values by applying a sigmoidal function; and (3) features from ESM2^47^, which is a recent protein language model trained on 15 billion parameters, leading to L×33 feature set, after normalizing the values using sigmoidal function.

Additionally, we extract a total of L×45 structure-based node features from the structure of the input monomer by either calculating various structural information directly from the 3D coordinates or by running the DSSP^48^ program. We describe them below.

##### Secondary structure and relative solvent accessibility

We use one-hot encoding of both 3-state and 8-state secondary structures (SS), leading to L×3 and L×8 feature sets, respectively. Additionally, we use one-hot encoding of 2-state relative solvent accessibility (RSA) by adopting an RSA cutoff of 50 (L×2 feature set), and use finer-grained RSA binning by discretizing into 8 bins as 0-30, 30-60, 60-90, 90-120, 120-150, 150-180, 180-210, and >210, represented by one-hot encoding (L×8 feature set). The rationale of using multiple discretization of SS (e.g., 3-state and 8-state) and RSA (e.g., 2-state and 8-state) is to combine multi-level information, and ablation studies guiding these decisions are presented in **Fig. 4**.

##### Local geometry

We use a total of L×11 feature set from the local geometries by calculating various planar and torsion angles of the polypeptide chain, including (1) the cosine angle between the consecutive residues of the C=O bond; (2) sine and cosine of the virtual bond and torsion angles formed between the consecutive C_α_ atoms; (3) normalized values of the backbone torsion angles.

##### Relative residue positioning

To capture the relative positional information for each residue, we extract two types of features: (1) for i^th^ residue, we use the inverse of i to capture the relative sequence position (L×1 feature set); and (2) we use the inverse of the Euclidean distance from the centroid of the input protein monomer to the C_α_ atom of the i^th^ residue to capture the spatial positioning of a residue relative to the overall structure (L×1 feature set).

##### Residue orientation

To define the orientation of each amino acid residue, we adopted features from a recent work^49^ including (1) the forward and reverse unit vectors in the directions of C_α_^(i+1)^ − C_α_^i^ and C_α_^(i−1)^ − C_α_^i^, respectively (L×6 feature set); and (2) the unit vector in the imputed direction of C_β_^i^ − C_α_^i^, computed by assuming tetrahedral geometries and normalization (L×3 feature set).

##### Residue virtual surface area

An amino acid residue can be perceived as a virtual convex hull constructed by its atoms. We calculate the virtual surface area of the convex hull and use its inverse as a feature (L×1 feature set).

##### Contact count

If the Euclidean distance between the C_β_ atoms of a residue pair is within a cutoff of 8Å, the two residues can be considered to be in contact. We calculate the contact-count by calculating the number of spatial neighbors of a given residue (i.e., residues which are in contact) and use the normalized number of contact count per residue as a feature (L×1 feature set).

#### Edge features

As the edge feature for the graph 𝒢 = (𝒱,ℰ), we calculate the ratio of the logarithmic sequential separation of two residues (i.e., logarithm of the absolute difference between the two residue indices) corresponding to two nodes in the graph and the Euclidean distance between them, defined as:

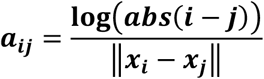

Here, the numerator captures how the two residues are separated in the primary sequence while the denominator captures their spatial interactions.

### Network architecture

We formulate the PPI site prediction into a graph node classification task and predict the probability of every residue in the input monomer to be a PPI site using a deep E(3) equivariant graph neural network. The network architecture consists of a stack of equivariant graph convolution layer (EGCL)^37^, performing a series of transformations of its input by updating the coordinate and node embeddings using the edge information and the coordinate and node embeddings from the previous layer. The EGCL operation attains equivariance primarily by changing the standard message passing 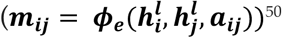 to equivariant message passing and by introducing coordinate updates in the graph neural network, as follows:

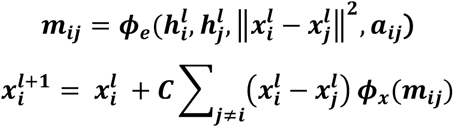

where ***a***_***ij***_ denotes edge features; 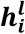, and 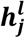 are node embeddings at layer ***l*** for nodes ***i*** and ***j***, respectively; ***ϕ***_***e***_, ***ϕ***_***h***_, and ***ϕ***_***x***_ are multilayer perceptrons (MLP) for edge, node, and coordinate operations, respectively. Equivariant message passing for an edge (***i***, ***j***) is attained by considering the squared distance between node ***i*** and 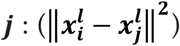 in the edge operation. The coordinate update for node ***i*** is obtained by the weighted sum of coordinate embedding difference from the previous layer, normalizing with a factor ***C*** = **1/(*M*** − **1)**, where ***M*** is the number of nodes in the graph. The weights for the sum are generated through the multilayer perceptron (MLP) of coordinate operation applied on the equivariant message passing (***m***_***ij***_) for each edge (***i***, ***j***).

Unlike off-the-shelf graph neural networks that aggregate messages only from the neighboring nodes, equivariant graph neural networks aggregate messages from the whole graph. Additionally, an attention embedding (***m***_***a***_) can be employed through an attention operation (***ϕ***_***a***_), which is a linear transformation on the aggregated message embedding (***m***_***i***_), followed by a sigmoidal non-linear transformation. The ‘attended’ aggregated message embedding is subsequently obtained through a scalar multiplication with the attention embedding. The node embedding is updated by applying an MLP (***ϕ***_***h***_) on the aggregated message and the node embeddings of the previous layer, as follows:

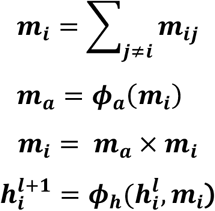

Finally, A linear transformation (***ϕ***_**0**_) is applied to squeeze the hidden dimension (***h***^***L***^) of the last EGCL, followed by a sigmoidal function to obtain the node-level classification (***p***_***i***_) for PPI site prediction:

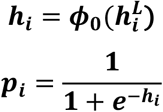

The network architecture of EquiPPIS consists of 10 layers of EGCL with a hidden dimension of 256, where the hyperparameters are chosen on an independent validation set, and empirical results guiding these decisions are presented in **Fig. 4**. EquiPPIS is implemented on Pytorch 1.12.0^51^ and Deep Graph Library (DGL) 0.9.0^52^. We use binary cross entropy loss function and cosine annealing scheduler from SGDR^53^, ADAM optimizer^54^ with a learning rate of 1e-4, and weight decay of 1e-16. The training process consists of at most 50 epochs on an NVIDIA A40 GPU. In addition to the full-fledged version of EquiPPIS, we train several baseline models on the same Train_335 dataset using the same set of input features and hyperparameters including off-the-shelf graph convolution network (GCN)^34^ and graph attention network (GAT)^44^, both implemented using the DGL^52^, as well as two variants of EquiPPIS: (1) ‘EquiPPIS invariant’, an invariant network with the coordinate updates of the equivariant graph convolution layers turned off; and (2) ‘EquiPPIS w/o attention’, an equivariant network with the attention operation turned off during equivariant message passing.

### Datasets, benchmarking, and performance evaluation

We use a combination of three widely used and publicly available benchmark datasets: Dset_186^6^, Dset_72^6^, and Dset_164^39^. Dset_72 is created based on protein-protein benchmark version 3.0^55^, Dset_186 is constructed through a six-step filtering process where 186 targets are collected from known protein complexes, and Dset_164 consists of 164 targets obtained from known heterodimers. While Dset_186, Dset_72, and Dset_164 are non-redundant data sets independently, the combined dataset is further reduced to 395 targets by filtering out redundant chains among the datasets. Following the same train-test split as GraphPPIS^9^, we use a train set (Train_335) having 10,374 and 55,992 interacting and noninteracting residues, respectively; and a test set (Test_60) having 2,075 and 11,069 interacting and noninteracting residues, respectively. In the Train_335 set, the average length of protein is ∼198 residues ranging from 44 to 869 residues. In the Test_60 set, the average length of protein is ∼219 residues ranging from 52 to 766 residues. Additionally, we adopt a dataset named Test_315 from the published work on GraphPPIS consisting of newly solved protein complexes that are non-redundant to the train set. We filter out 42 targets from the Test_315 set by discarding protein chains having more than 25% pairwise sequence identity with our test set and create an independent validation set (Validation_42) to perform feature ablation and hyperparameter selection.

During prediction, we use both experimentally-solved structures as well as on AlphaFold2-predicted structural models as input. We run AlphaFold2 with default parameter settings by locally installing the officially released version^29^ to generate five predicted structural models and then select the model with the highest pLDDT confidence score. For target 4cdgA that failed during the MSA generation stage of the AlphaFold2 pipeline, we run Colabfold^56^ that uses MMSeqs2^57^ for MSA generation and subsequently employs AlphaFold2 protocol for structure prediction.

EquiPPIS is compared against both sequence-based (PSIVER^6^, ProNA2020^40^, SCRIBER^41^, DLPred^42^, and DELPHI^8^) and structure-aware (DeepPPIS^7^, SPPIDER^10^, MaSIF-site^43^, and GraphPPIS^9^) PPI site prediction methods. PSIVER employs a Naïve Bayes classifier along with kernel density estimation by utilizing sequence-based features. ProNA2020 combines homology modeling with a neural network for residue-level PPI site prediction. SCRIBER employs two layers of logistic regression, where the first layer utilizes sequence-based features while the second layer combines the output from the first layer for the final prediction. DLPred employs a simplified long-short-term memory model for PPI site prediction. DELPHI uses an ensemble of convolutional and recurrent neural networks architectures with a large feature set. DeepPPISP combines local contextual features with global features and employs convolutional neural networks to predict PPI sites. SPPIDER leverages support vector machine, neural network, and linear discriminant analysis with an extensive feature search and extraction process. MaSIF-site predicts PPI sites by learning protein structural fingerprints through geometric deep learning. GraphPPIS employs structure-aware deep residual neural networks for PPI site prediction.

For benchmarking and performance assessment, we use standard performance evaluation metrics including accuracy, precision, recall, F1-score (F1), and Matthews correlation coefficient (MCC), defined as:

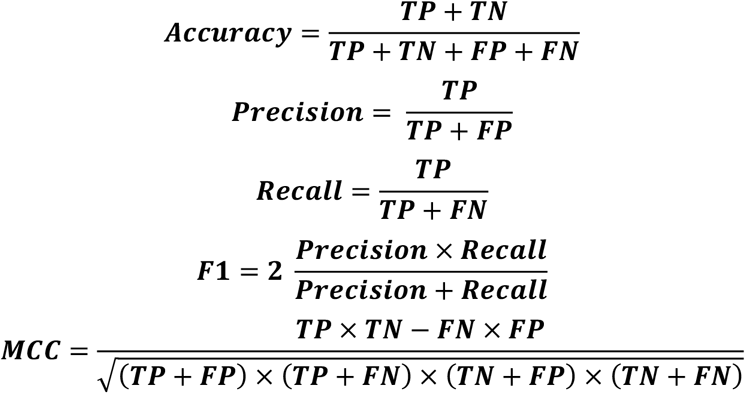

where, TP denotes the number of true PPI site residues that are correctly predicted, FP denotes the number of non-PPI site residues that are incorrectly predicted to be in PPI sites, TN denotes the number of non-PPI site residues that are correctly predicted, and FN denotes the number of PPI site residues that are incorrectly predicted as non-PPI site. We additionally use area under the receiver operating characteristic curve (ROC-AUC) and area under the precision-recall curve (PR-AUC) for performance evaluation.

## Data availability

The raw data used in this study, including the datasets for train, test and validation are collected from publicly available sources and freely available at https://github.com/biomed-AI/GraphPPIS.

## Code availability

An open-source software implementation of EquiPPIS, licensed under the GNU General Public License v3, is freely available at https://github.com/Bhattacharya-Lab/EquiPPIS.

## Acknowledgements

This work was partially supported by the National Institute of General Medical Sciences [R35GM138146 to D.B.] and the National Science Foundation [DBI2208679 to D.B.].

